# COVID-19 and Rheumatoid Arthritis share myeloid pathogenic and resolving pathways

**DOI:** 10.1101/2020.07.26.221572

**Authors:** Lucy MacDonald, Thomas D. Otto, Aziza Elmesmari, Barbara Tolusso, Domenico Somma, Charles McSharry, Elisa Gremese, Iain B. McInnes, Stefano Alivernini, Mariola Kurowska-Stolarska

## Abstract

**Background:** We recently delineated the functional biology of pathogenic and inflammation resolving synovial tissue macrophage clusters in rheumatoid arthritis (RA). Whilst RA is not a viral respiratory syndrome, it represents a pro-inflammatory cytokine-driven chronic articular condition often accompanied by cardiovascular and lung pathologies. We hypothesised that functionally equivalent macrophage clusters in the lung might govern inflammation and resolution of COVID-19 pneumonitis.

**Methods:** To provide insight into the targetable functions of COVID-19 bronchoalveolar lavage (BALF) macrophage clusters, a comparative analysis of BALF macrophage single cell transcriptomics (scRNA-seq) with synovial tissue (ST) macrophage scRNA-seq and functional biology was performed. The function of shared BALF and ST MerTK inflammation-resolving pathway was confirmed with inhibitor in primary macrophage-synovial fibroblast co-cultures. **Results**. Distinct BALF FCN^**pos**^ and FCN^**pos**^SPP1^**pos**^ macrophage clusters emerging in severe COVID-19 patients were closely related to ST CD48^**high**^S100A12^**pos**^ and CD48^**pos**^SPP1^**pos**^ clusters driving synovitis in active RA. They shared transcriptomic profile and pathogenic mechanisms. Healthy lung resident alveolar FABP4^pos^ macrophages shared a regulatory transcriptomic profile, including TAM (Tyro, Axl, MerTK) receptors pathway with synovial tissue TREM2^pos^ macrophages that govern RA remission. This pathway was substantially altered in BALF macrophages of severe COVID-19. In vitro dexamethasone inhibited tissue inflammation via macrophages’ MerTK function.

**Conclusion:** Pathogenesis and resolution of COVID-19 pneumonitis and RA synovitis might be driven by similar macrophage clusters and pathways. The MerTK-dependent anti-inflammatory mechanisms of dexamethasone, and the homeostatic function of TAM pathways that maintain RA in remission advocate the therapeutic MerTK agonism to ameliorate the cytokine storm and pneumonitis of severe COVID-19.

## Introduction

The clinical, social and economic disruption caused by SARS-CoV-2 infection and its unpredictable progression to COVID-19 and Acute Respiratory Distress Syndrome (ARDS) represents a global emergency. The severity of COVID-19 is attributable to immune dysregulation, abnormal blood clotting and tissue disruption, particularly implicating pro-inflammatory innate immunity ^1-3^. Interstitial lung disease and alveolitis is a co-morbidity of rheumatoid arthritis (RA) ^4^, in which articular inflammation and disease remission are driven by distinct synovial tissue macrophage clusters ^5^. Despite suggestive early data ^2 6-10^ there is a knowledge gap of how myeloid cell pathways mechanistically regulate severity or contribute to the resolution of COVID-19, hindering development of effective treatments ^11-14^. Single-cell profiling and fate-mapping indicate spatial and functional macrophage heterogeneity that maintains lung homeostasis ^15-19^. The alveolar macrophages (AM) expressing Fatty Acid Binding Protein 4 (FABP4) localise to the alveolar epithelial surface and recycle surfactants with type-2 epithelial cells to maintain compliance and efficient gas exchange ^20^. This function is compromised in severe COVID-19 ^21^. Recent scRNAseq analysis ^22^ of bronchoalveolar lavage fluid (BALF) from 6 severe COVID-19 patients found abnormally low numbers of resident alveolar macrophages and an increase in two macrophage clusters which share expression of ficolin-1 (FCN1), and can be distinguished by their relative expression of osteopontin (SPP1): FCN^**pos**^ and FCN^**pos**^SPP1^**pos**^ clusters. Their roles in the pathogenesis of severe COVID-19 have yet to be established. We recently identified similar macrophage diversity in synovial tissue (ST) of healthy donors and patients with active or remission rheumatoid arthritis (RA) ^5^. Whilst RA is not a viral respiratory syndrome, it represents a pro-inflammatory cytokine-driven chronic articular condition often accompanied by cardiovascular and lung pathologies ^4 23^. We ^24^ and others ^25^ have reported that SARS-CoV-2 infection is associated with emergence of polyarthritis or flares of synovitis in RA patient in sustained disease remission, suggesting potential shared pathogenic mechanisms of COVID-19 and RA. RA immunopathogenesis and therapeutic targets ^23^ are well understood and thus might be informative for COVID-19 therapeutic strategies. For example, dexamethasone is a drug of choice for flares of arthritis and recent efficacy studies of dexamethasone ^26^ highlights opportunities to understand alveolar macrophage-based mechanism of resolution of COVID-19. We recently delineated the functional biology of pathogenic and inflammation resolving synovial tissue macrophage clusters and their roles in pathology or resolution of RA synovitis ^5^. Thus, we raised the hypothesis that functionally equivalent macrophage clusters in the lung govern inflammation and resolution of COVID-19 pneumonitis. We propose that this perspective can offer new insight into the role of innate immunity in the immunopathogenesis and resolution of COVID-19.

## Methods

### Dataset Acquisition

Our analysis strategy involved the comparison and integration of 10X Genomics single cell RNA sequencing data of myeloid cells from bronchoalveolar lavage fluid (BALF) of Liao et al (2020)^22^ and from synovial tissue (ST) of Alivernini et al (2020)^5^. Sequencing data from BALF was acquired in the format of CellRanger output from GEO under the accession number GSE145926. Synovial tissue macrophage data was acquired from EMBL-EBI using the accession numbers: *E-MTAB-8322*.

### Quality checks, filtering and clustering of the scRNAseq datasets (Supplementary Figure.1)

The Seurat package (3.1.2) in R was used to create an object (CreateSeuratObject). Cell filtering involved removal of cells with less than 500 expressed genes (subset, subset=nFeatures_RNA > 500). We also set thresholds for level of gene expression, including expression of mitochondrial genes (percent.mt). This allows for exclusion of doublets and dying cells. The data was normalized using Seurat’s NormalizeData function. For the analysis of myeloid cells only, cells were filtered for expression of CD14, MARCO, LYZ with the subset function. The top 2000 variable genes were then identified for all samples, using the FindVariableFeatures function. *Integration*: Sample integration of each dataset was performed following the Seurat vignette, integrating all genes that are common between samples of each condition, using the functions: FindIntegrationAnchors, and IntegrateData (features.to.integrate to normalize all genes common to samples). *Individual Clustering and Dimensional Reduction*: UMAP based on PCA cell embeddings were generated by Seurat for each dataset using RNA assay. For visualization of ST data, the first 12 principal components (PCs) are used (RunUMAP). The same PCs were used in determination of the k-nearest neighbours for each cell during SNN graph construction before clustering at a chosen resolution of 0.5 (FindNeighbors, FindClusters) and identifying the populations described in Alivernini et al (2020)^5^. In comparison, the first 50 principle components (PCs) and a resolution of 0.8 were used for visualization and clustering of the BALF data as described in Laio et al (2020)^22^ methods. The populations described by Laio et al (2020)^22^ were identified by merging clusters based on expression of FCN1, SPP1 and FABP4. The relative proportion of clusters between conditions was explored by exporting tables of percentage of cells in each cluster for each sample and performing statistical analyses using Prism (8.4.2). Kruskal-Wallis test with Dunn’s correction for multiple comparisons was used. *Differential Expression Analysis*. In order to identify cluster markers, the Seurat function FindAllMarkers was used with each dataset individually. The “test.use” function was used to determine genes differentially expressed between clusters within each dataset using MAST. As recommended, for DE comparison the non-batch normalized counts were used. For identification of cluster markers, we specify that any markers identified must be expressed by at least 40% of cells in the cluster (‘min.pct’ parameter 0.4). We use the default values for all other parameters. Genes are considered significantly DE if the adjusted p-value (< 0.05) by Bonferroni Correction and multiple test correction (multiple by number of tests/clusters).

### Dataset Comparison

#### Integration

Following individual clustering analysis, dataset integration was performed using the same method as described above, integrating 8993 common genes between ST and BALF data. These “integrated” batch-corrected values were then set as the default assay and the gene expression values are scaled before running principal component analysis. *Clustering and Dimensional Reduction*. To prevent bias in clustering, we reduced cell number of BALF data to 32139 random cells to match ST data. The data was re-scaled (ScaleData) before the first 30 principal components (PCs) were used for UMAP visualization. Cells were coloured by their original identity to illustrate how clusters of each dataset overlay. *Comparative analyses*. A count matrix with the average expression of all common genes by each cluster was generated (AverageExpression) before hierarchical clustering (dist, hclust) and plotting a dendrogram. Principal component analysis (PCA) was also performed on this pseudobulk expression matrix (prcomp). These initial analyses allowed for the identification of similar clusters between BALF and ST data. Shared marker genes between similar clusters were identified by comparing marker gene lists of each cluster using Venny (2.1) To visualise pseudobulk expression of the overlapping marker genes, the pheatmap (1.0.12) package was used to visualize shared genes as a heatmap using a custom script. Comparison of the pseudobulk of expression marker genes unique and common to all clusters similar between datasets was performed by Pearson correlation (ggscatter, add = “reg.line”, conf.int = TRUE, cor.coef = TRUE, cor.method = “pearson”). These were performed based on the 916, 721 and 986 unique and common genes of BALF FCN1^pos^/ST CD48^high^S100A12^pos^, BALF FCN^pos^SPP1^pos^/ST CD48^pos^SPP1^pos^ and BALF FABP4^pos^/ST TREM2^pos^, respectively.

### Pathway Analysis

To investigate the functions of shared and unique genes for BALF and ST clusters obtained in the analysis above, the pathway analysis using the Reactome^27^ (database v. 73) was performed. Genes shared between the clusters that showed the same direction of change in both datasets were submitted to the Reactome. Differentially expressed genes (DE) were split by direction of log2FC gene (upregulated log2FC > 0 and downregulated log2FC < 0) before submission to the Reactome. Identified pathways were filtered for p-value<0.05, FDR (False Discovery Rate)<0.1, confidence levels with minimum entities found>4, and minimum entities ratio>0.01. Plots were made using Jupyter notebook 6.0.3, pandas 0.23.4 and seaborn 0.10.1.

### Co-culture of macrophages with synovial fibroblasts

#### Direct co-culture system

CD14^**pos**^ cells were isolated from peripheral blood mononuclear cells (PBMC) using CD14 micro-beads (Miltenyi Biotec) with the autoMACS Pro Separator according to the manufacturer’s protocol. Sorted monocytes were differentiated to monocyte-derived macrophages in complete medium containing macrophage colony-stimulating factor (M-CSF). Briefly, cells were plated in a 6-well culture plate at a density of 1.5×10^6^ per well in 3ml of RPMI 1640 medium supplemented with 10% FCS, penicillin/streptomycin (100 U/ml), and 2mM Glutamine in the presence of M-CSF 50ng/ml (PeproTech, UK). Cells were replenished on day 3 with fresh medium containing M-CSF (50ng/ml). On day 6, cells were pre-treated (16h) with LPS (1ng/ml) to model inflammatory macrophages or dexamethasone (1μM) to model regulatory macrophages or both, in the presence of absence of MerTK inhibitor UNC1062^28^ (Aobious; 250nM). After extensive washing, cells were co-cultured in a flat bottom 96-wells plate at a density of at 2×10^3^ per well with RA synovial fibroblasts (^5^ that seeded 24h before co-culture, at 37°C in 5%CO2 controlled environment for different periods of time. Prior to co-culture, macrophages and synovial fibroblasts were labelled with CellTrace™-Far Red (5 μM, Life Technologies) and CellTrace™-Violet (5 μM, Life Technologies), respectively according to the manufacturer’s protocol. After 24 & 48 hours of co-culture, culture supernatants were collected for assay of mediators, while macrophages and synovial fibroblasts were sorted based on positivity for Far Red (macrophages) or CellTrace™ Violet (synovial fibroblasts) into RLT buffer (Qiagen) containing 1% β-mercaptoethanol using FACS-ARIA III, and expression set of MMPs and IL-6 in both macrophages and synovial fibroblasts were analysed by ultrasensitive qPCR. The MMP Luminex panel (PPX-05/ PROCARTAPLEX MMP1, MMP2, MMP3, MMP9 and MMP13 plex) and IL-6 Elisa (both from Life Technologies) was performed on co-cultures supernatants.

#### qPCR for MMPs and IL-6

RNA from synovial fibroblasts was isolated using the RNEasy micro-kit (Qiagen), and cDNA was prepared using a High Capacity cDNA Reverse Transcription Kit (Thermo Fisher Scientific). TaqMan mRNA primer/probe assays and TaQman Gene Expression master mixes (both from Life Technologies) were used for semi-quantitative determination of genes of interest. Data is presented as relative value (*i*) 2^-ΔCT^ where ΔCt=Cycle threshold for 18S (housekeeping) minus Ct for gene of interest, or (*ii*) fold-change, where ΔCt for selected control condition =1 or 100%.

We used the following primers/ probe TaqMan assays:

Hs00174131_m1/ IL-6,

Hs00899658_m1/MMP1,

Hs00968305_m1/MMP3,

#### Statistical evaluation of the frequency of BALF and ST macrophage clusters and co-culture experiments

Two-way Anova (Tukey’s correction) or One-way Anova (Dunn’s correction) to evaluate the differences between clusters across all groups, or Mann-Whitney between two groups. To evaluate the influence of distinct MoM phenotypes on FLS functions, One-way Anova with Dunn’s correction for multiple comparisons, or paired t-test was used if 2 groups were compared. The name of statistical test and p-values and range or exact n numbers are provided in each Figure.

## Results

### COVID-19 BALF FCN1^pos^ and FCN1^pos^SPP1^pos^ macrophage clusters are similar to the CD48^high^S100A12^pos^ and CD48^pos^SPP1^pos^ clusters that drive synovitis

In COVID-19 BALF, Liao et al ^22^ identified 4 major clusters of macrophages characterised by a combination of expression of SPP1, FCN1 and FABP4. Importantly they found that expansion of FCN1^**pos**^ and FCN1^**pos**^SPP1^**pos**^ clusters was indicative of COVID-19 severity (**Fig.1a-c**). Among the 9 phenotypically distinct clusters of ST macrophages that differed in distribution between health and RA, the CD48^**high**^S100A12^**pos**^ and CD48^**pos**^SPP1^**pos**^ clusters were expanded in RA patients with active disease ^5^ (**Fig.1d-f**). To test the relationship between COVID-19 BALF and RA ST macrophage clusters, we integrated the macrophage scRNAseq datasets from both COVID-19 BALF ^22^ **(Fig.1a-c)** and RA ST ^5^ (**Fig.1d-f**). The dimensional reduction of integrated macrophage datasets illustrated by Uniform Manifold Approximation and Projection (UMAP) visualise potentially overlapping and tissue-specific macrophage clusters (**Fig.1g-h**). To investigate the relationship between these macrophage clusters of synovitis and pneumonitis, a hierarchical clustering analysis (**Fig.1i**) and principal component analysis (**Fig.1j**) was performed on the matrix of the average expression of each ST and BALF cluster of all 8902 genes that were common to both datasets. The hierarchical clustering dendrogram illustrating the relationship between the clusters by branch-point (split) and branch-length (distance) revealed that the macrophage clusters were separated predominantly by their function *e*.*g*. embryonic-origin/homeostasis vs monocyte-derived/inflammation, and not by tissue source. The first branch-point separated 4 tissue-resident clusters of healthy lung (FABP4^**pos**^ and FABP4^**low**^SPP1^**pos**^) and synovial lining (TREM2^**high**^ and TREM2^**low**^) macrophages from the other BALF and ST macrophage clusters that were likely of monocytic origin ^29^. Similarly, the leading principal component that accounts for 54% of the integrated dataset variability largely separated the clusters found in homeostatic conditions from those in diseased lung and synovium (**Fig.1j**). The split-points and distances in the second branch of hierarchical clustering indicate that the proinflammatory clusters (*i*.*e*. FCN^**pos**^ and FCN^**pos**^SPP1^**pos**^) which dominate in severe COVID-19 share transcriptomic profiles with pro-inflammatory clusters (*i*.*e*. CD48^**high**^S100A12^**pos**^ and CD48^**pos**^SPP1^**pos**^) present in active RA, respectively. Pearson correlation analysis between BALF FCN1^**pos**^ and ST CD48^**high**^S100A12^**pos**^, and between BALF FCN1^**pos**^SPP1^**pos**^ and ST CD48^**pos**^SPP1^**pos**^ confirmed the similarities (r=0.56, p=2.2^-16^ and r=0.65, p=2.2^-16^ for respective BALF/ST pairs). The most striking pathogenic similarities were between the COVID-19 pneumonitis FCN1^**pos**^ and the active RA synovitis CD48^**high**^S100A12^**pos**^ clusters. They share 238 top marker-genes (**Fig.1k**) that include up-regulation of inflammatory and prothrombotic pathways. These consisted of IFN pathway (*e*.*g. IFITM2, IFITM3 and ISG15*), inflammation-triggering alarmins (*S100A8/9/12A)*, B-cell activation factors (*e*.*g*. BAFF-*TNFSF15*B), promotors of IL-1β and TNF production (*e*.*g. CARD16* and *LITAF*), prothrombotic factors (*e*.*g. FGL2*) and integrins mediating cell migration and adhesion (*e*.*g. ITGB2* and *ITGX*). They also share receptor expression profiles *e*.*g*. TNFR2 (*TNFSF1*), G-CSF receptor (*CSFR3*) and the complement receptor (*C5aR*) that may increasingly render them susceptible to pro-inflammatory mediators to escalate inflammation (**Fig.1l** and **Supplementary Fig.2**). The hierarchical analysis and comparison of cluster markers also indicate a shared transcriptional profile of COVID-19 FCN1^**pos**^SPP1^**pos**^ and synovitis CD48^**pos**^SPP1^**pos**^ clusters consisting of 86 common marker-genes including *SPP1*, which have broad pro-inflammatory functions ^30^ (**Fig.1m-n** and **Supplementary Fig.3**). These transcriptomic profiles strongly suggest that BALF FCN^**pos**^ and FCN^**pos**^SPP1^**pos**^ macrophage clusters of COVID-19 patients shared pathogenic mechanisms with ST CD48^**high**^S100A12^**pos**^ and CD48^**pos**^SPP1^**pos**^ of active RA.

**Figure 1.**
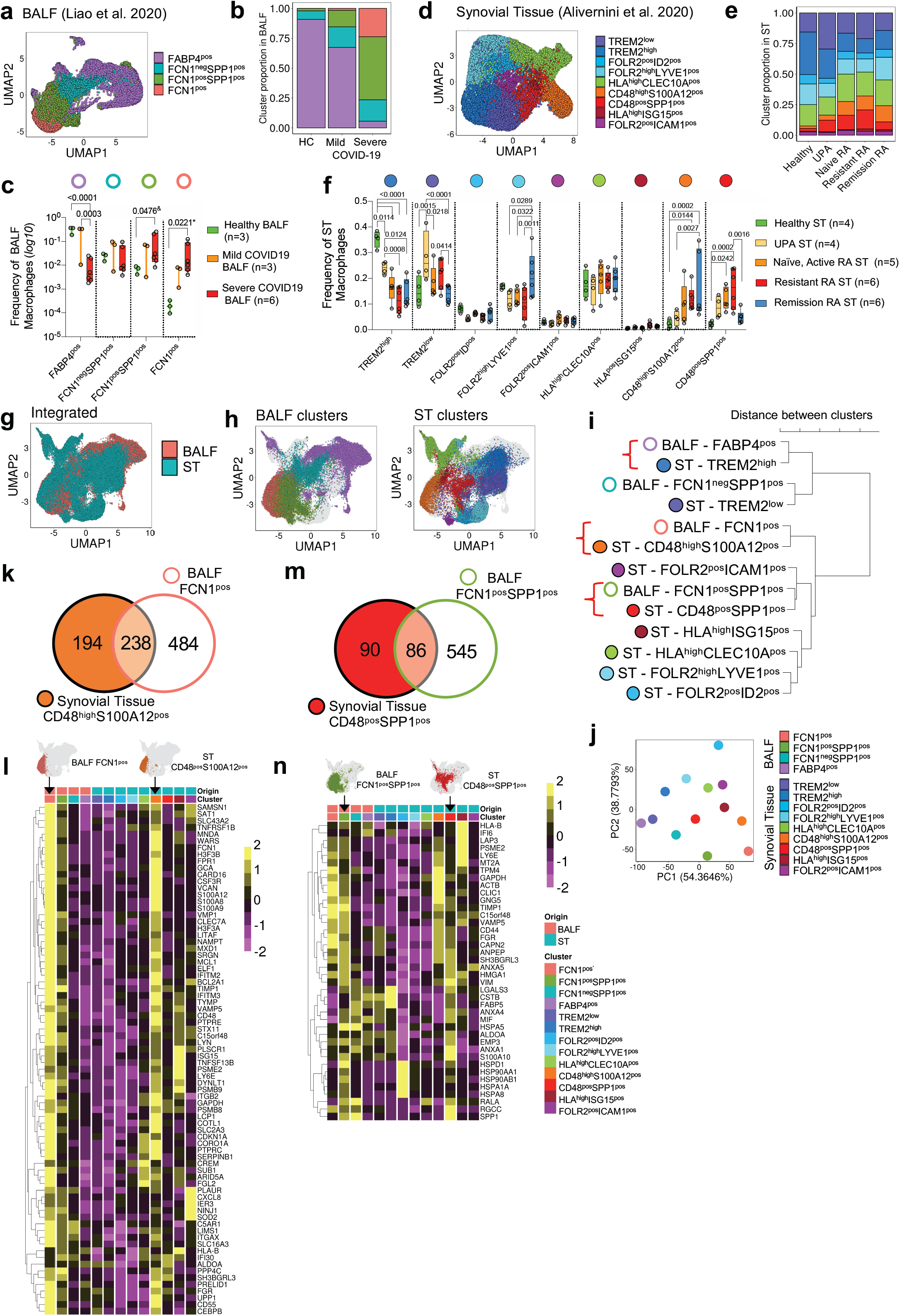
COVID-19 BALF FCN1^pos^ and FCN1^pos^SPP1^pos^ macrophage clusters are transcriptionally similar to the CD48^high^S100A12^pos^ and CD48^pos^SPP1^pos^ clusters of synovial tissue macrophages from active RA. (**a**) UMAP of 52,380 BALF macrophages^22^. Each dot represents one cell and is colored by cluster identity. (**b**) Stacked-bar and (**c**) Box-and-whisker plots illustrating BALF macrophage cluster proportions between conditions. (**d**) UMAP of 32,139 synovial tissue (ST) macrophages^5^. Each dot represents one cell and is colored by cluster identity. (**e**) Stacked-bar and (**f**) Box-and-whisker plots illustrating ST macrophage cluster proportions between conditions. Two-way Anova (Tukey’s correction) or one-way Anova (Dunn’s correction) (*) to evaluate the differences between clusters across all groups or Mann-Whitney (^&^) between two groups. Exact P-values are on the graphs. (**g**-**h**) UMAP of integrated ST and BALF macrophages (**g**) and colored according to their ST and BALF cluster identification (**h**). (**i-j**) Dendrogram of hierarchical clustering analysis (**i**) and PCA (**j**) of integrated pseudo-bulk gene expression (average expression in each cluster) of ST and BALF clusters. (**k**) Venn diagram illustrating numbers of unique and shared marker-genes of ST CD48^**high**^S100A12^**pos**^ and BALF FCN1^pos^ clusters. p-value <0.05 after Bonferroni correction. (**l**) Heatmap illustrating scaled, pseudo-bulk gene expression of shared upregulated marker-genes (highlighted in k) by BALF and ST clusters. (**m**) Venn diagram illustrating numbers of unique and shared marker-genes of ST CD48^**pos**^SPP1^**pos**^ and BALF FCN1^**pos**^SPP1^**pos**^ clusters generated as in (k). (**n**) Heatmap illustrating scaled, pseudo-bulk gene expression of shared upregulated marker-genes (highlighted in m) by ST and BALF clusters. HC, healthy control; UPA, undifferentiated arthritis; Naïve, RA naïve to any treatment.

### Tissue resident alveolar FABP4^pos^ macrophages share regulatory transcriptomic profile, including TAM pathway with synovial tissue lining layer TREM2^pos^ macrophages

While the innate immune pathways that contribute to the resolution of COVID-19 are yet to be characterised, recently uncovered innate mechanisms reinstating tissue homeostasis in RA ^5^ might shed new light on potential resolution mechanisms of COVID-19. In ST of RA patients in sustained disease remission we identified novel macrophage clusters with protective and inflammation-resolving properties. One of these clusters, TREM2^**pos**^ grouped tightly with the FABP4^**pos**^ alveolar macrophage cluster in unsupervised hierarchical analysis (**Fig1i**). Comparison of gene expression profiles confirmed that they shared transcriptomics (r=0.83, p=2.2^-16^ in a Pearson correlation of 986 of TREM2^**pos**^ and FABP4^**pos**^ unique and common cluster-markers) that might indicate analogous homeostatic functions (**Fig.2a-b**). In the synovium, the TREM2^**pos**^ macrophages form a lining-layer producing and recycling lubricant synovial fluid which facilitates joint movement ^31^ while in lung alveoli, the FABP4^**pos**^ macrophages form part of the alveolar barrier producing and recycling surfactants to maintain airway patency and facilitate gas exchange. The BALF FABP4^**pos**^ and the ST TREM2^**high**^ clusters share 170 markers (**Fig.2a-b** and **Supplementary Fig.4**). These include the complement pathways (*e*.*g. C1q*, that facilitates uptake of apoptotic bodies), high expression of genes of retinoic acid production (*e*.*g. ALDH1A1* and *RBP4*) driving T-reg differentiation ^32^, and B7-related co-inhibitory molecule *VSIG4* which inhibits T-effector cells ^33^, together suggesting a primary role of this cluster in governing lung immunity. While the functional contribution of FABP4^**pos**^ macrophages to the resolution of SARS-CoV-2 infection is yet to be established, we found that their counterpart TREM2^**pos**^ ST macrophage clusters produced inflammation-resolving lipid mediators and induce a repair phenotype in tissue stromal cells that maintain disease remission. These homeostatic responses are driven by MerTK, a member of the immunosuppressive tyrosine kinase receptor TAM family (namely Tyro, Axl and MerTK respectively) ^5^. TAM receptors and their ligands GAS6 or PROS1 form a homeostatic brake on inflammation and autoimmunity ^34-36^. In addition, PROS1 is an essential inhibitor of blood coagulation preventing thrombosis ^37^. Lung-resident FABP4 macrophages uniquely express Axl rather than MerTK. Their Axl is constitutively ligated to GAS6 ^38 39^ and is key in preventing exacerbated inflammation *e*.*g*. during influenza virus infection ^38 39^. Analysis of the Liao COVID-19 dataset ^22^ identified profoundly altered macrophage expression of TAM receptors and their ligands in the lung (**Fig.2c**) that might explain the inadequate regulation of tissue hyper-inflammatory response in severe COVID-19. Alveolar macrophages from healthy lungs show high expression levels of Axl and PROS1, and these were markedly reduced in severe COVID-19 patients. GAS6 and MerTK are not expressed by resident AMs; instead they were increasingly expressed in infiltrating FCN^**pos**^ and FCN^**pos**^SPP1^**pos**^ macrophage clusters, suggesting an inflammation-triggered attempt to counterbalance pathogenic responses. However, the reduced PROS1, which is the preferred activating-ligand for MerTK ^36 40^, and fewer resident homeostatic AMs in severe COVID-19 might enable the unrestricted proinflammatory cytokine production by locally differentiated MerTK-positive FCN^**pos**^ and FCN^**pos**^SPP1^**pos**^ macrophages, while the reduced PROS1 contribute to increased local coagulopathy. In addition, the markedly reduced Axl expression in the remaining AMs of severe COVID-19 may additionally impair their ability to control inflammatory response and their homeostatic gas exchange functions.

**Figure 2.**
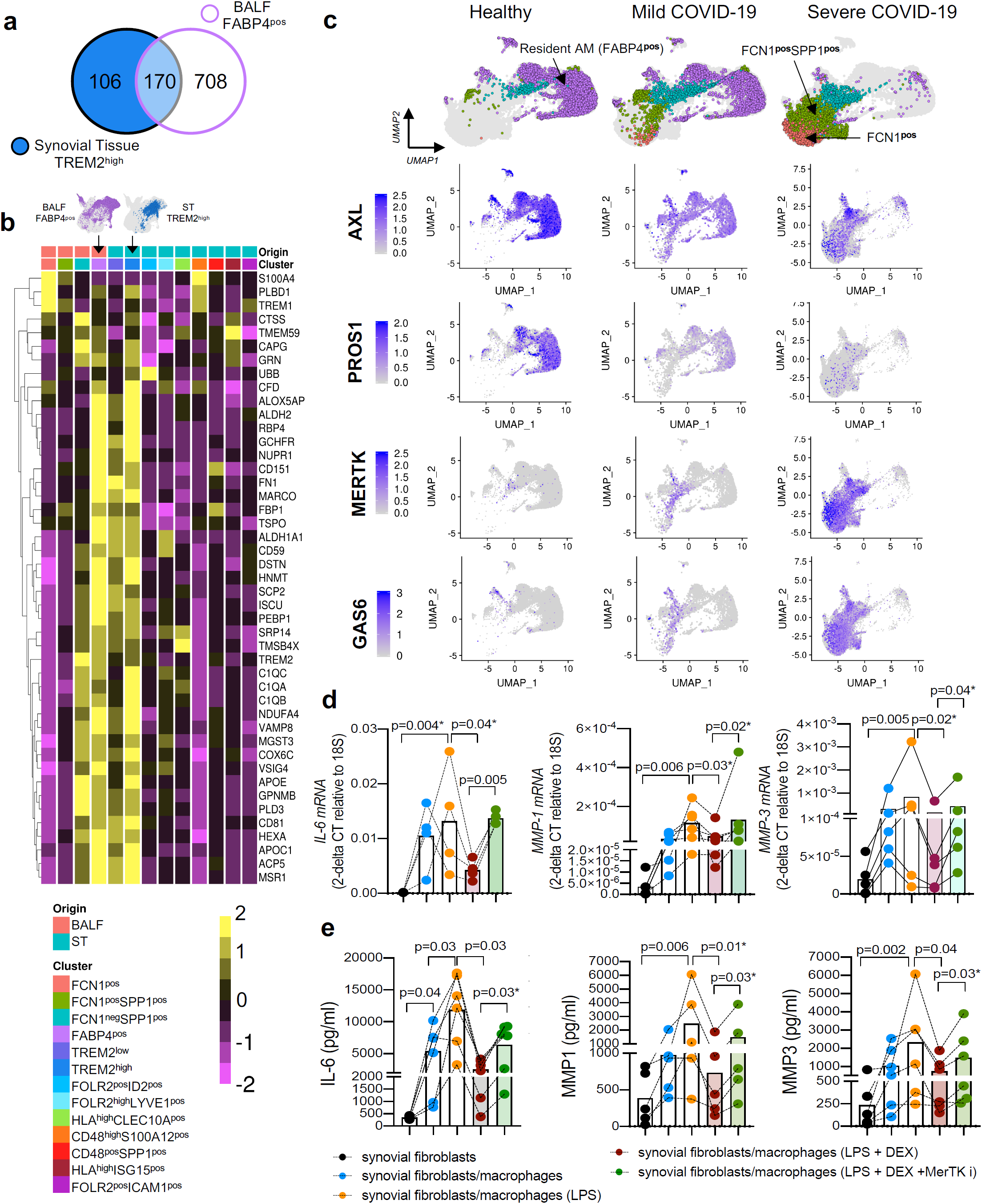
COVID-19 BALF FABP4^pos^ and RA synovial TREM2^pos^ macrophages share transcriptomic profiles and regulatory TAM receptor pathways. (**a**) Venn diagram illustrating numbers of unique and shared marker-genes of ST TREM2^**high**^ and BALF FABP4^**pos**^ macrophage clusters. Marker-genes were identified prior to integration of datasets and were calculated using MAST, setting minimum percentage of cells in clusters expressing each marker to 40%. Genes considered differentially expressed when p<0.05 after Bonferroni correction. (**b**) Heatmap illustrating scaled, pseudo-bulk expression of shared upregulated marker-genes from ST and BALF clusters indicated in (a). (**c**) Split UMAP plots comparing BALF macrophage clusters in health, and in mild and severe COVID-19 illustrating changes in expression of TAM receptors Axl and MerTK with their respective preferred ligands PROS1 and GAS6. Intensity of purple indicates expression level. (**d**) mRNA levels of *IL6, MMP1* and *MMP3* in synovial fibroblasts (SF: FACS-sorted from co-cultures with different macrophage phenotypes). (**e**) Supernatant concentrations of IL-6, MMP-1 and MMP-3 from co-cultured SF with different macrophage phenotypes: (**d-e**) MerTK inhibition of LPS+dexamethasone pre-treated macrophages enhanced SF expression and co-culture concentrations of pro-inflammatory mediators. Data presented as dot-plots of individual SF donors (n=5 in 3 independent experiments) with bar indicating the mean. One-way Anova, followed by Dunn’s correction, or two-sided paired t-test of two groups (*). P-values are on the graphs. SF, synovial fibroblasts; ST, synovial tissue; BALF, bronchoalveolar lavage fluid; Dex, Dexamethasone.

### Dexamethasone inhibits tissue inflammation via MerTK function in macrophages

Preliminary findings from the RECOVERY trial (no_NCT04381936) of the corticosteroid dexamethasone in COVID-19 showed reduced mortality of patients requiring mechanical ventilation and has been approved for treatment of hospitalised COVID-19 patients ^26^. Understanding the underlying mechanisms of its efficacy may improve targeted therapy for COVID-19. Dexamethasone strongly induces macrophage expression of immunosuppressive MerTK and GAS6 ^41^, thus restoration of their homeostatic functions might contribute to the decreased mortality. Since dexamethasone is a treatment of choice for acute flares of synovitis in RA patients ^42^ we used an *ex-vivo* synovitis model to test mechanisms of efficacy that might also be relevant for COVID-19. We co-cultured primary synovial fibroblasts from RA patients with different phenotypes of human monocyte-derived macrophages. These included macrophages pre-treated with (***i***) TLR4 ligand LPS (*to model pro-inflammatory macrophages*), or with (***ii***) TLR4 ligand LPS plus dexamethasone (*to model MerTK-positive homeostatic tissue macrophages*), or with (***iii***) LPS/dexamethasone plus MerTK inhibitor to test the specificity of MerTK involvement. Co-culture of fibroblasts with (***i***) pro-inflammatory macrophages induced a pathogenic phenotype expressing IL-6 and MMPs in the co-cultured stromal cells. Induction of this pathogenic fibroblast phenotype was inhibited in co-culture with (***ii***) the same pro-inflammatory macrophages pre-treated with dexamethasone. This anti-inflammatory effect of dexamethasone was attenuated by blocking macrophage MerTK leading to increased expression/production of IL-6 and MMPs by stromal cells (**Fig.2d-e**). Together this suggests that at least in part, dexamethasone inhibits tissue inflammation via inducing MerTK function in macrophages.

## Discussion

*In summary*, this comparative single-cell transcriptomic analysis of myeloid cells from COVID-19 pneumonitis and myeloid cells from chronic synovitis and remission RA suggests that their pathogenesis and resolution might be driven by similar myeloid cell clusters/pathways (**Fig.3**). Promising data from COVID-19 trials testing drugs already used for the treatment of RA *e*.*g*. anti-IL-6R ^12^, anti-GM-CSF ^43 44^ and dexamethasone ^26^ further support the concept of common pathogenic and resolution mechanisms that, if appropriately targeted, might prevent/resolve severe COVID-19 pneumonitis.

**Figure 3.**
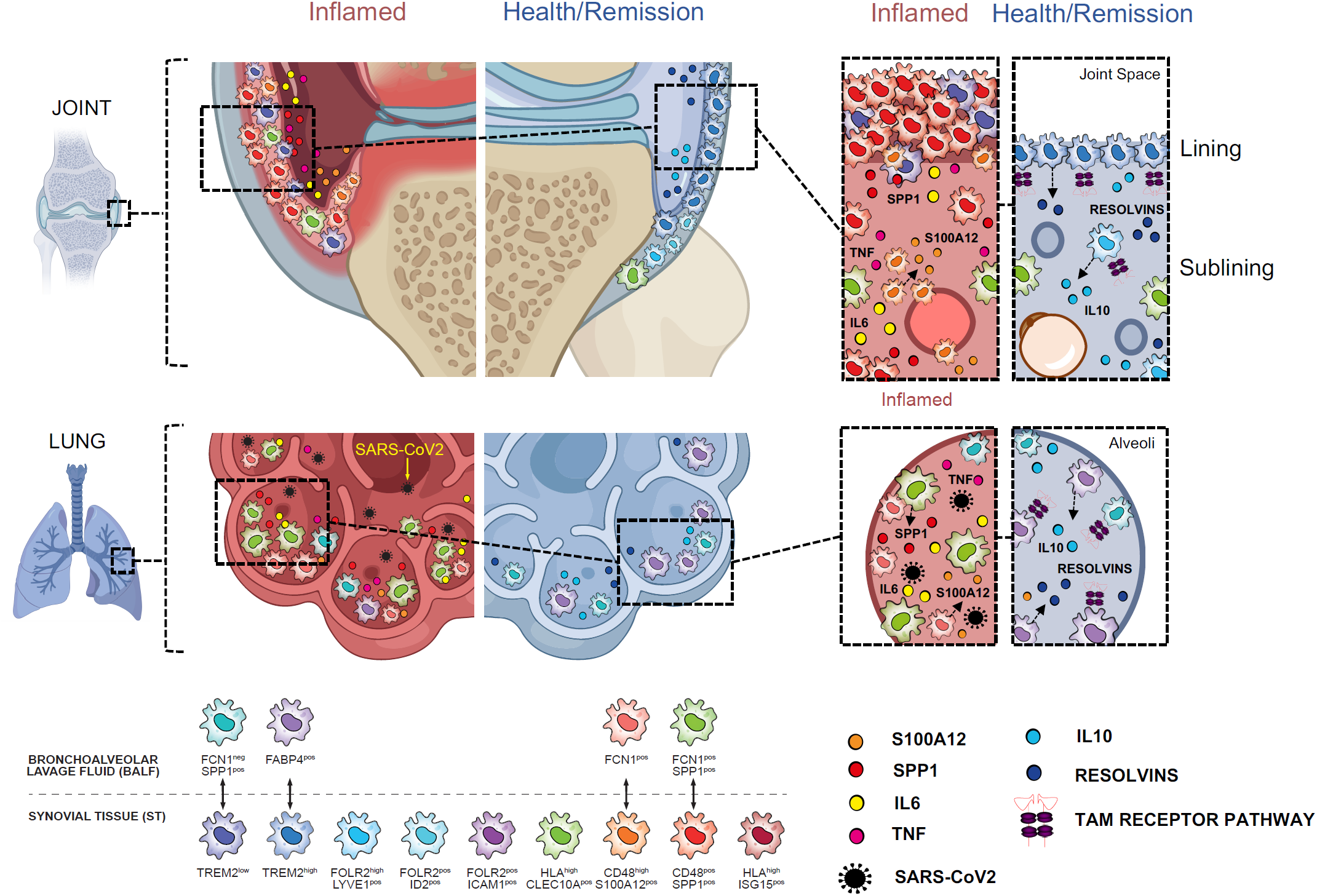
COVID-19 and Rheumatoid Arthritis share pathogenic and resolving myeloid pathways. Bronchoalveolar and synovial tissue macrophages are heterogenous. Healthy lung contains FABP4^**high**^ alveolar macrophages that maintain efficient gas exchange, and a smaller FABP4^**low**^SPP1^**pos**^ cluster with yet unknown function. During SARS-CoV-2 infection, distinct FCN1^**pos**^ and FCN^**pos**^SPP1^**pos**^ macrophage clusters emerge in alveoli. In healthy joints, predominant TREM2^**high**^ and TREM2^**low**^ with a contribution of FOLR2^**pos**^LYVE1^**pos**^ macrophage clusters form the synovial lining layer while the sublining layer is populated by FOLR2^**pos**^LYVE1^**pos**^, FOLR2^**pos**^ID2^**pos**^, FOLR2^**pos**^ICAM1^**pos**^ macrophage clusters, and myeloid CLEC10a^**pos**^MHCII^**high**^ dendritic cells. During RA, new CD48^**high**^S100A12^**pos**^ and CD48^**pos**^SPP1^**pos**^ synovial tissue (ST) macrophage clusters emerge and these are the main producers of pro-inflammatory mediators and induce pathogenic changes in adjacent stromal tissue. In COVID-19, the BALF FCN1^**pos**^ and FCN^**pos**^SPP1^**pos**^ macrophage clusters transcriptionally resemble the pathogenic ST CD48^**high**^S100A12^**pos**^ and CD48^**pos**^SPP1^**pos**^ clusters, suggesting that they might similarly mediate pathogenic pneumonitis contributing to severity of COVID-19. Disease remission in RA is associated with the regulatory functions of TREM2^**pos**^ and FOLR2^**pos**^LYVE1^**pos**^ macrophage clusters that produce inflammation-resolving mediators and induce homeostatic repair in stromal cells, mediated by MerTK and its ligand GAS6. MerTK is a member of the tyrosine kinase receptor TAM family (Tyro, Axl, MerTK), and ligating TAM receptors by ligands GAS6/PROS1 forms a homeostatic brake on inflammation. Healthy alveolar FABP4^**pos**^ macrophages transcriptionally resemble arthritis resolving TREM2^**pos**^ macrophages, suggesting potentially similar homeostatic mechanisms. These alveolar FABP4^**pos**^ macrophages constitutively express Axl and the MerTK ligand PROS1. Expression of these is substantially decreased in severe COVID-19, and this may contribute to unrestricted proinflammatory cytokine production by infiltrating MerTK expressing FCN^**pos**^ and FCN^**pos**^SPP1^**pos**^ creating a cytokine storm. ST, synovial tissue; BALF, bronchoalveolar lavage fluid; RA, rheumatoid arthritis.

The transcriptomic profiles strongly suggest that BALF FCN^**pos**^ and FCN^**pos**^SPP1^**pos**^ macrophage clusters of COVID-19 patients shared pathogenic pathways with ST CD48^**high**^S100A12^**pos**^ and CD48^**pos**^SPP1^**pos**^ of active RA. While the precise functions of these BALF clusters remain unclear, recently dissected functional biology of their *ex-vivo* synovial counterparts (*i*.*e*. CD48^**high**^S100A12^**pos**^ and CD48^**pos**^SPP1^**pos**^) revealed they were the main producers of pathogenic TNF, IL-6, IL-1β, alarmins and chemokines in the synovium, and they also induced contact-dependent emergence of a pathogenic cluster of tissue stromal cells producing tissue-destructive factors (*e*.*g*. MMPs) ^5^. Thus, this comparative analysis together with high concentrations of pro-inflammatory mediators in BALF of severe COVID-19 patients ^22^ reveals FCN^**pos**^ and FCN^**pos**^SPP1^**pos**^ macrophage subpopulations as potential perpetrators of pulmonary hyper-inflammatory and tissue-damaging innate immune responses towards SARS-CoV-2 infection. Further functional studies of the interaction between these pathogenic macrophage subpopulations and lung epithelial and stromal cells could clarify mechanisms of pneumonitis and potential repurposing of current anti-inflammatory biologics for COVID-19 ^11 45^.

We uncovered that tissue resident alveolar FABP4^**pos**^ macrophages and ST lining layer TREM2^**pos**^ macrophages share transcriptomic profiles and regulatory TAM (Tyro, Axl, MerTK) receptor pathways that were profoundly repressed in severe COVID-19. TAM pathways are widely shared between resident macrophages of many tissues ^41 46^ and have clinical consequences when deregulated. For example, a low proportion of MerTK-expressing ST macrophages in RA patients in sustained remission is predictive of joint flare after drug withdrawal ^5^. Similarly, experimental arthritis is greatly exacerbated in MerTK-deficient mouse ^47^ and triple TAM receptor-deficient mice spontaneously develop autoimmunity ^48^. Thus, deregulation of the TAM pathway could explain the COVID-19 related hyper-inflammatory response, increased lung blood clotting, and breach of self-tolerance *e*.*g*. production of anti-phospholipid antibodies ^49^. The MerTK-dependent anti-inflammatory mechanisms of dexamethasone, and the homeostatic function of TAM pathways that maintain RA patients in disease remission ^5^ strongly recommend therapeutic MerTK agonism to ameliorate the cytokine storm and pneumonitis of severe COVID-19 ^37^.

## Key messages

### What is already known about this subject?

COVID-19 clinical trials testing anti-inflammatory drugs used for the treatment of RA *e*.*g*. anti-IL-6R, anti-GM-CSF and dexamethasone support the concept of some common COVID-19 and RA pathogenic and resolution mechanisms. Underlying shared innate cellular and molecular mechanisms are unknown and if dissected could provide novel therapeutic targets to resolve severe COVID-19 pneumonitis.

### What does this study add?

- Lung BALF macrophage clusters in severe COVID-19 (FCN^**pos**^ and FCN^**pos**^SPP1^**pos**^) share pathogenic mechanisms with pro-inflammatory macrophage clusters that drive synovitis (CD48^**high**^S100A12^**pos**^ and CD48^**pos**^SPP1^**pos**^).
- Healthy alveolar macrophages (FABP4^pos^) and healthy and remission synovial tissue lining layer macrophages (TREM2^pos^) share transcriptomic profiles, including the regulatory TAM (Tyro, Axl, MerTK) receptor pathway. This pathway is altered in BALF macrophages in severe COVID-19.
- Dexamethasone inhibits inflammation via macrophages’ MerTK function.

### How might this impact on clinical practice or future developments?

- COVID-19 pneumonitis is driven by macrophage clusters similar to those that drive chronic arthritis, and importantly, that resolution of COVID-19 might be accelerated by engaging similar macrophage-resolving pathways as found in remission arthritis, providing putative candidate inflammation-resolving therapeutic pathways (e.g MerTK) that might prevent/resolve severe COVID-19 pneumonitis.

## Acknowledgments

This work was supported by the Research into Inflammatory Arthritis Centre *Versus Arthritis* UK (grant nos. 20298 and 22072) to L.M., T.D.O., I.B.M. and M.K-S.; Università Cattolica del Sacro Cuore (no. R4124500654) to S.A.; Ricerca Finalizzata Ministero della Salute (no. GR-2018-12366992, to S.A.); Versus Arthritis UK Program (grant no. 21802, to M.K-S); Versus Arthritis UK (grant no. 22272, to M.K-S.); Wellcome Trust (no. 204820/Z/16/Z, to T.D.O. and M.K-S.); and BPF-Medical Research Trust (to C.M.).

## Author Contribution

**L.M**. analysed scRNAseq data, interpreted all the results and wrote the manuscript. **T.D.O**. and **M.K.-S** supervised all computational analysis in the study. **M.K.-S**. and **S.A** conceived and oversaw the project, interpreted all the results and wrote the manuscript with feedback from all authors. **S.A**. and **B.T**. performed all synovial biopsies. **A.E**. performed a direct cocultures of MoM with FLS. **D.S**. performed pathway analysis **I.B.M** and **E.G**. assisted with running the project and help with result interpretation.

## Ethical approval statement

The study was approved by the Ethic Committee of the Fondazione Policlinico Universitario A. Gemelli IRCCS – Università Cattolica del Sacro Cuore, Rome, Italy.

## Patient and public involvement statement

Patients or the public were not involved in the design, conduct, reporting or dissemination of our research.

## Figure legends

**Supplementary Figure 1.**
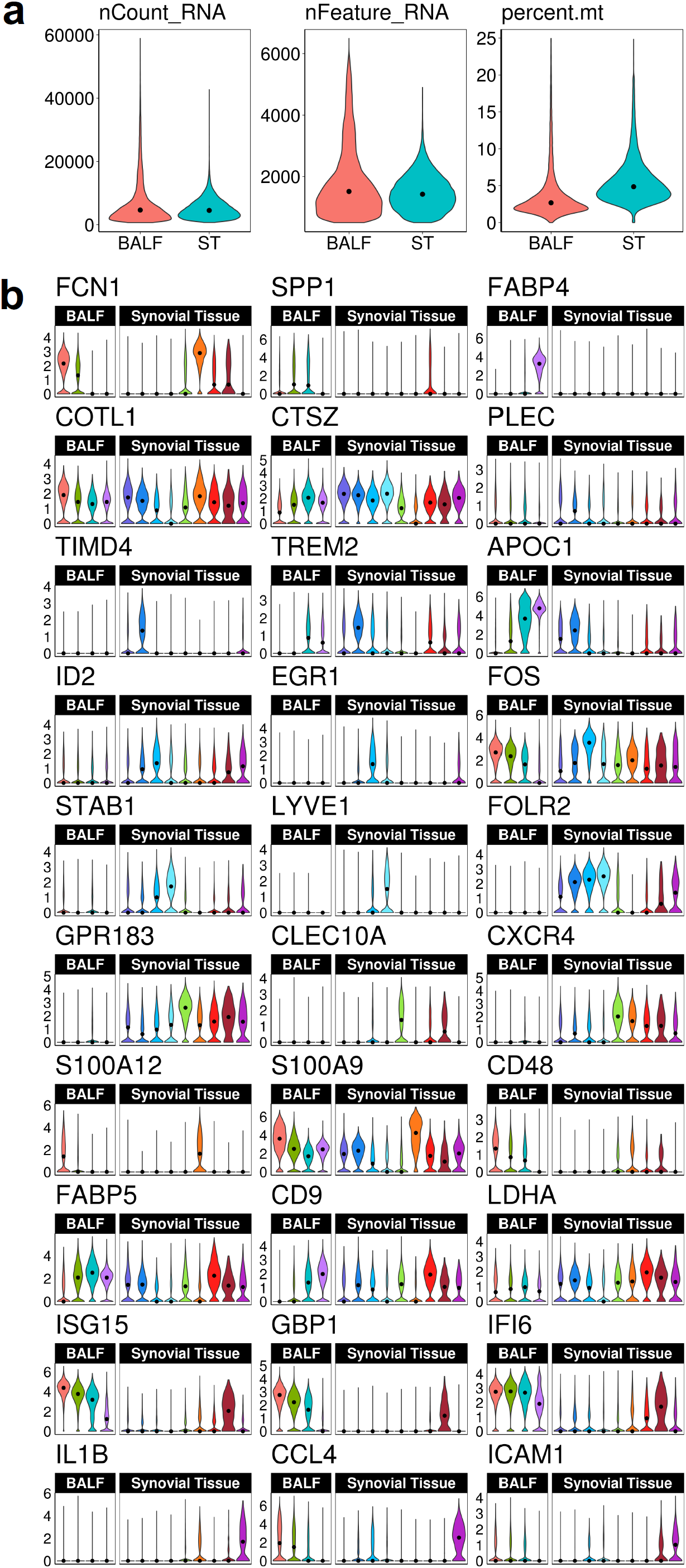
Quality checks of scRNAseq data sets. **(a)** Violin plots comparing the average number of counts (nCount_RNA), genes (nFeature_RNA) and percentage of mitochondrial genes (percent.mt) per cell per dataset. The black dot represents the median value per dataset. (**b**) Violin plots illustrating marker gene expression across all BALF and synovial tissue (ST) clusters. X-axis represents distinct clusters with violins colored by cluster identity. Y-axis represents distribution of log-normalized expression values of associated cluster marker with median value of each cluster marked by black dot.

**Supplementary Figure 2.**
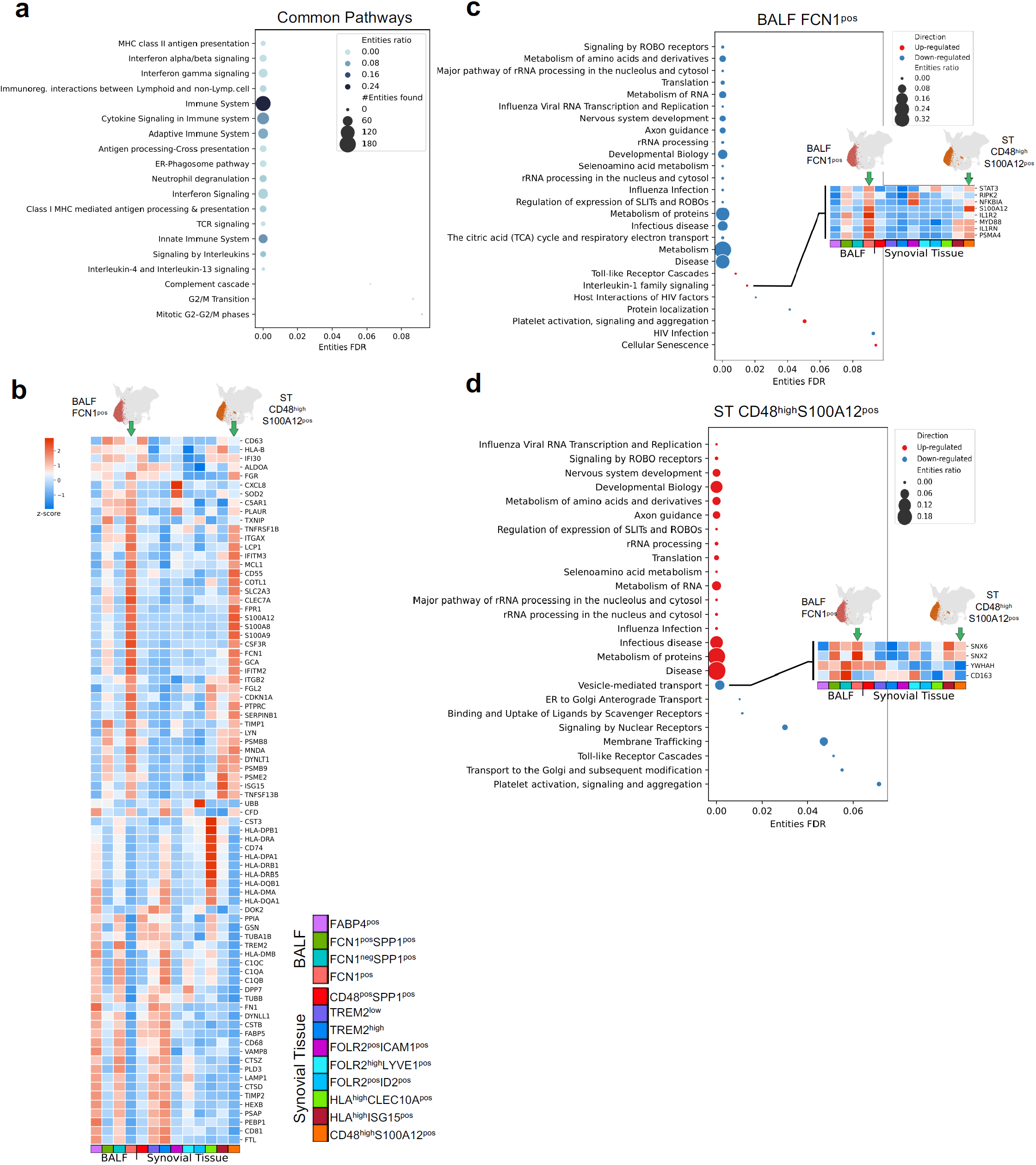
Shared and unique pathways of BALF FCN1^pos^ and ST CD48^high^S100A12^pos^ macrophages. (**a**) Common pathways of BALF FCN1^**pos**^ and ST CD48^**high**^S100A12^**pos**^ macrophages. Each dot represents a significantly enriched pathway. Pathway names are indicated on the y-axis and the false discovery rate (FDR) is on the x-axis. Number of entities and entities-ratio are indicated by dot-size and darker colour, respectively. (**b**) Heatmap illustrating z-score of pseudo-bulk expression of genes in the pathways indicated in (a). Arrows point to BALF FCN1^**pos**^ and ST CD48^**high**^S100A12^**pos**^. (**c**) Pathways which differ between BALF FCN1^**pos**^ and ST CD48^**high**^S100A12^**pos**^ clusters. (**d**) Pathways which differ between ST CD48^**high**^S100A12^**pos**^ and BALF FCN1^**pos**^ clusters. (c-d) Each dot represents a significantly enriched pathway. Pathway names are indicated on y-axis while FDR on x-axes. Entities-ratio is indicated by dot-size, and colours indicate pathways upregulated/downregulated. Small heatmaps illustrate z-score of pseudo-bulk expression of genes in the indicated pathway. Arrows point to BALF FCN1^**pos**^ and ST CD48^**high**^S100A12^**pos**^ clusters. Pathway analysis performed with Reactome (database v. 73).

**Supplementary Figure 3.**
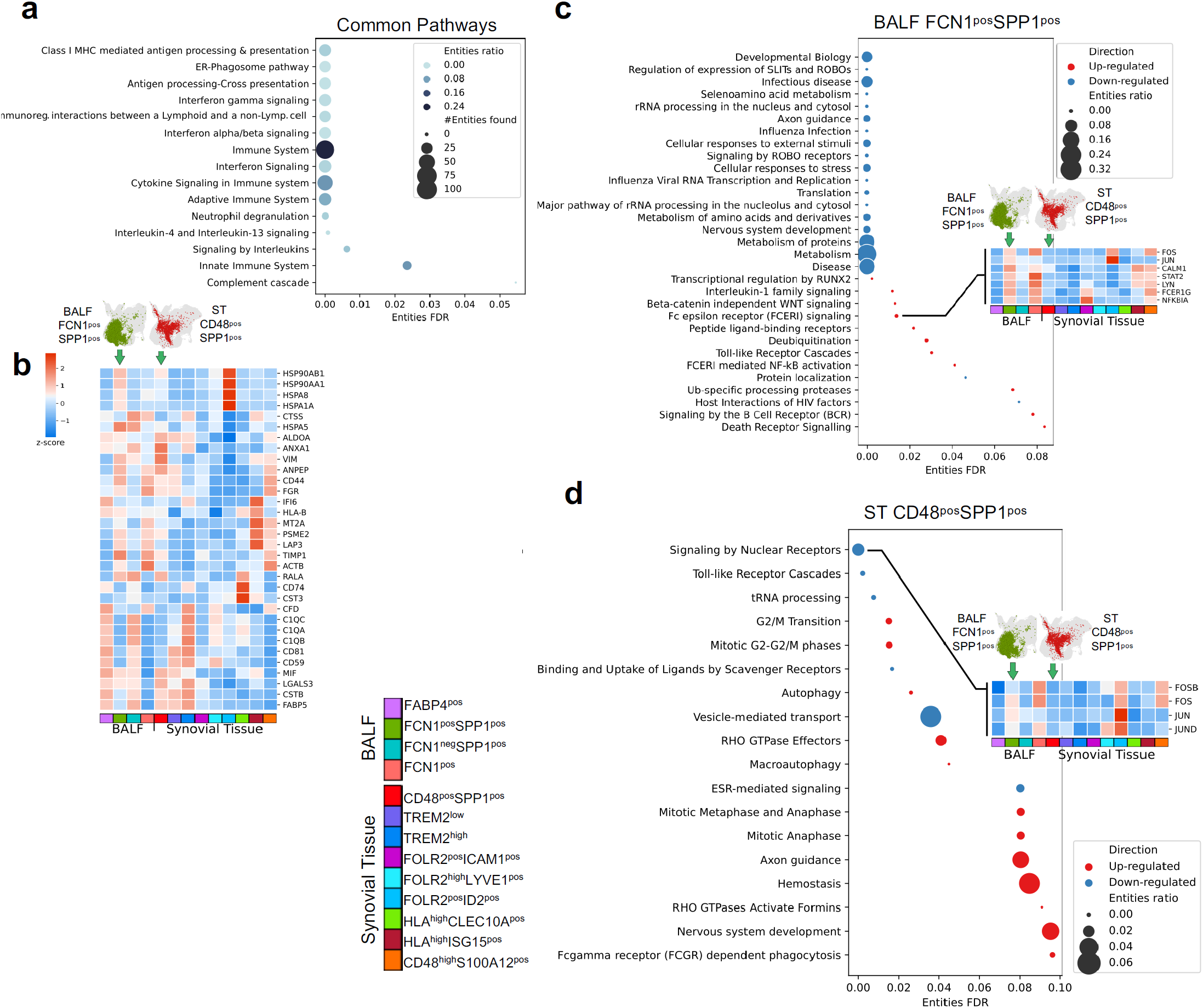
Shared and unique pathways of BALF FCN1^pos^SPP1^pos^ and ST CD48^pos^SPP1^pos^ macrophages. (**a**) Common pathways of BALF FCN1^**pos**^SPP1^**pos**^ and ST CD48^**pos**^SPP1^**pos**^ macrophages. Each dot represents a significantly enriched pathway. Pathway names are indicated on y-axis and false discovery rate (FDR) on x-axes. Number of entities and entities ratio are indicated with dot size and darker colour, respectively. (**b**) Heatmap illustrating z-score of pseudo-bulk expression of genes in the pathways indicated in (a). Arrows point to BALF FCN1^**pos**^SPP1^**pos**^ and ST CD48^**pos**^SPP1^**pos**^. (**c**) Pathways which differ between BALF FCN1^**pos**^SPP1^**pos**^ and ST CD48^**pos**^SPP1^**pos**^ clusters. (**d**) Pathways which differ between ST CD48^**pos**^SPP1^**pos**^ and BALF FCN1^**pos**^SPP1^**pos**^ clusters. (c-d) |Each dot represents a significantly enriched pathway. Pathway names are indicated on y-axis and FDR on x-axes. Entities-ratio is indicated with dot-size, and colours indicate pathways upregulated/downregulated. Small heatmaps illustrate z-score of pseudo-bulk expression of genes in the indicated pathway. Arrows point to BALF FCN1^**pos**^SPP1^**pos**^ and ST CD48^**pos**^SPP1^**pos**^ clusters. Pathway analysis performed with Reactome (database v.73).

**Supplementary Figure 4.**
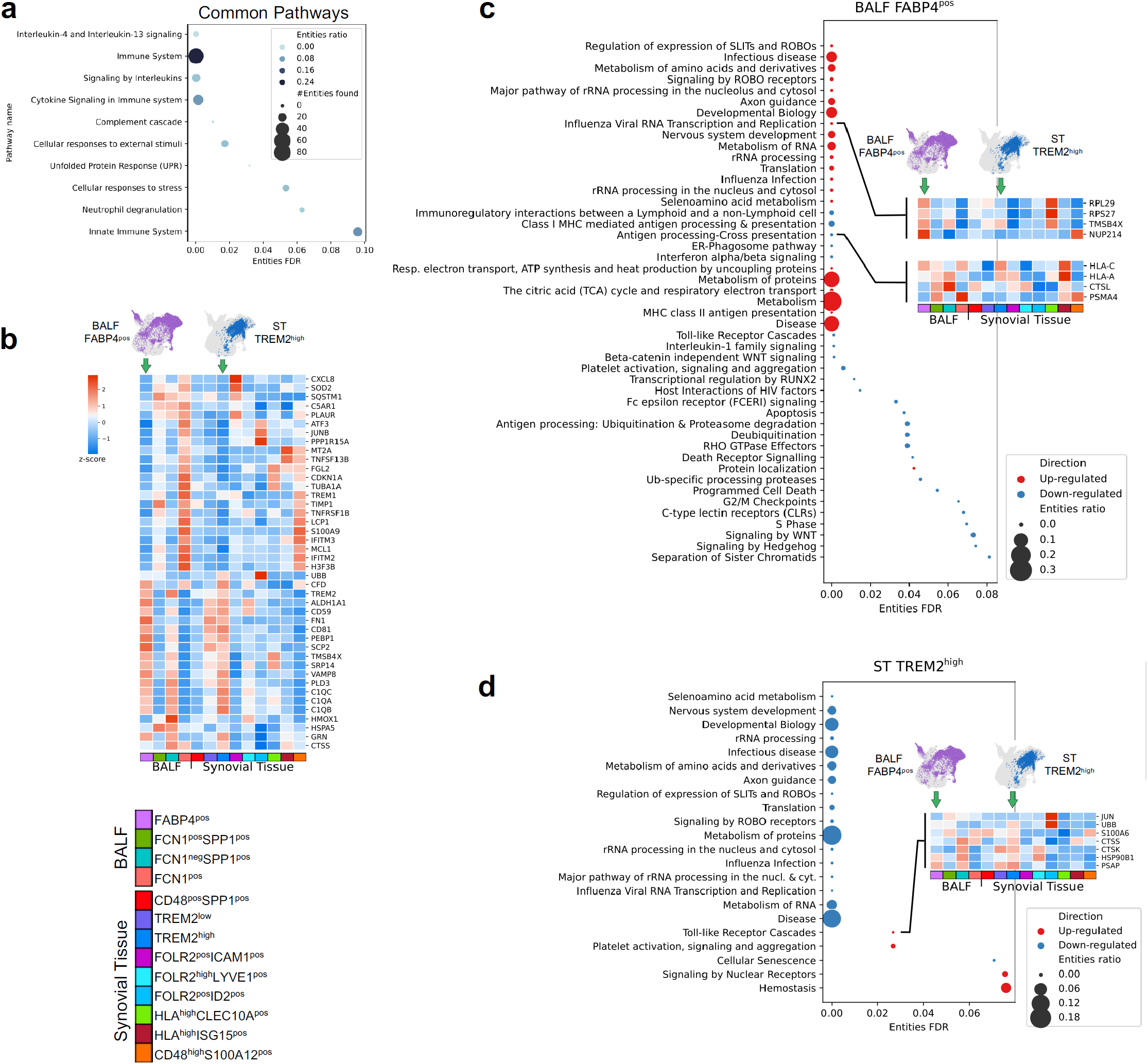
Shared and unique pathways of BALF FABP4^pos^ and ST TREM2^high^ macrophages. (**a**) Common pathways of BALF FABP4^**pos**^ and ST TREM2^**high**^ macrophages. Each dot represents a significantly enriched pathway. Pathway names are indicated on y-axis and false discovery rate (FDR) on x-axes. Number of entities and entities-ratio are indicated by dot-size and darker colour, respectively. (**b**) Heatmap illustrating z-score of pseudo-bulk expression of genes in the pathways indicated in (a). Arrows point to BALF FABP4^**pos**^ and TREM2^**high**^. (**c**) Pathways which differ between BALF FABP4^**pos**^ and ST TREM2^**high**^ clusters. (**d**) Pathways which differ between ST TREM2^**high**^ and BALF FABP4^**pos**^ cluster. (c-d) Each dot represents a significantly enriched pathway. Pathway names are indicated on y-axis and FDR on x-axes. Entities-ratio is indicated by dot-size, and colours indicate pathways upregulated/downregulated. Small heatmaps illustrate z-score of pseudo-bulk expression of genes in the indicated pathway. Arrows point to BALF FABP4^**pos**^ and ST TREM2^**high**^ clusters. Pathway analysis performed with Reactome (database v.73).

